# Sex-specific genome-wide association study in glioma identifies new risk locus at 3p21.31 in females, and finds sex-differences in risk at 8q24.21

**DOI:** 10.1101/229112

**Authors:** Quinn T. Ostrom, Ben Kinnersley, Margaret R. Wrensch, Jeanette E. Eckel-Passow, Georgina Armstrong, Terri Rice, Yanwen Chen, John K. Wiencke, Lucie S. McCoy, Helen M. Hansen, Christopher I. Amos, Jonine L. Bernstein, Elizabeth B. Claus, Dora Il’yasova, Christoffer Johansen, Daniel H. Lachance, Rose K. Lai, Ryan T. Merrell, Sara H. Olson, Siegel Sadetzki, Joellen M. Schildkraut, Sanjay Shete, Joshua B. Rubin, Justin D. Lathia, Michael E. Berens, Ulrika Andersson, Preetha Rajaraman, Stephen J. Chanock, Martha S. Linet, Zhaoming Wang, Martha S. Linet, Zhaoming Wang, Meredith Yeager, on behalf of the GliomaScan consortium, Richard S. Houlston, Robert B. Jenkins, Beatrice Melin, Melissa L. Bondy, Jill. S. Barnholtz-Sloan

## Abstract

Incidence of glioma is approximately 50% higher in males. Previous analyses have examined exposures related to sex hormones in women as potential protective factors for these tumors, with inconsistent results. Previous glioma genome-wide association studies (GWAS) have not stratified by sex. Potential sex-specific genetic effects were assessed in autosomal SNPs and sex chromosome variants for all glioma, GBM and non-GBM patients using data from four previous glioma GWAS. Datasets were analyzed using sex-stratified logistic regression models and combined using meta-analysis. There were 4,831 male cases, 5,216 male controls, 3,206 female cases and 5,470 female controls. A significant association was detected at rs11979158 (7p11.2) in males only. Association at rs55705857 (8q24.21) was stronger in females than in males. A large region on 3p21.31 was identified with significant association in females only. The identified differences in effect of risk variants do not fully explain the observed incidence difference in glioma by sex.

## Introduction

Glioma is the most common type of primary malignant brain tumor in the United States (US), with an average annual age-adjusted incidence rate of 6.0/100,000 [1]. Glioma can be broadly classified into glioblastoma (GBM, 61.9% of gliomas in adults 18+ in the US) and lower-grade glioma (non-GBM glioma, 24.2% of adult gliomas) with tumors such as ependymoma (6.3%), unclassified malignant gliomas (5.1%), and pilocytic astrocytoma (1.9%) making up the majority of other cases [1]. Many environmental exposures have been investigated as sources of glioma risk, but the only validated risk factors for these tumors are ionizing radiation (which increases risk), and history of allergies or other atopic disease (which decreases risk) [2]. These tumors are significantly more common in people of European ancestry, in males and in older adults [1]. The contribution of common low-penetrance SNPs to the heritability of sporadic glioma in persons with no documented family history is estimated to be ∼25% [3]. A recent glioma genome-wide association study (GWAS) meta-analysis validated 12 previously reported risk loci [4], and identified 13 new risk loci. These 25 loci in total are estimated to account for ∼30% of heritable glioma risk. This suggests that there are both undiscovered environmental risk (which accounts for ∼75% of incidence variance) and genetic risk factors (accounting for ∼70% of heritable risk) [3,4].

Population-based studies consistently demonstrate that incidence of gliomas varies significantly by sex. Most glioma histologies occur with a 30-50% higher incidence in males, and this male preponderance of glial tumors increases with age in adult glioma (**Figure 1**) [1]. Several studies have attempted to estimate the influence of lifetime estrogen and progestogen exposure on glioma risk in women [5,6]. Results of these analyses have been mixed, and it is not possible to conclusively determine the impact of hormone exposure on glioma risk. Male predominance in incidence occurs broadly across multiple cancer types and is also evident in cancers that occur in pre-pubertal children and in post-menopausal adults [7,8]. Together these observations suggest that other mechanisms in addition to acute sex hormone actions must be identified to account for the magnitude of sex difference in glioma incidence.

**Figure 1.**
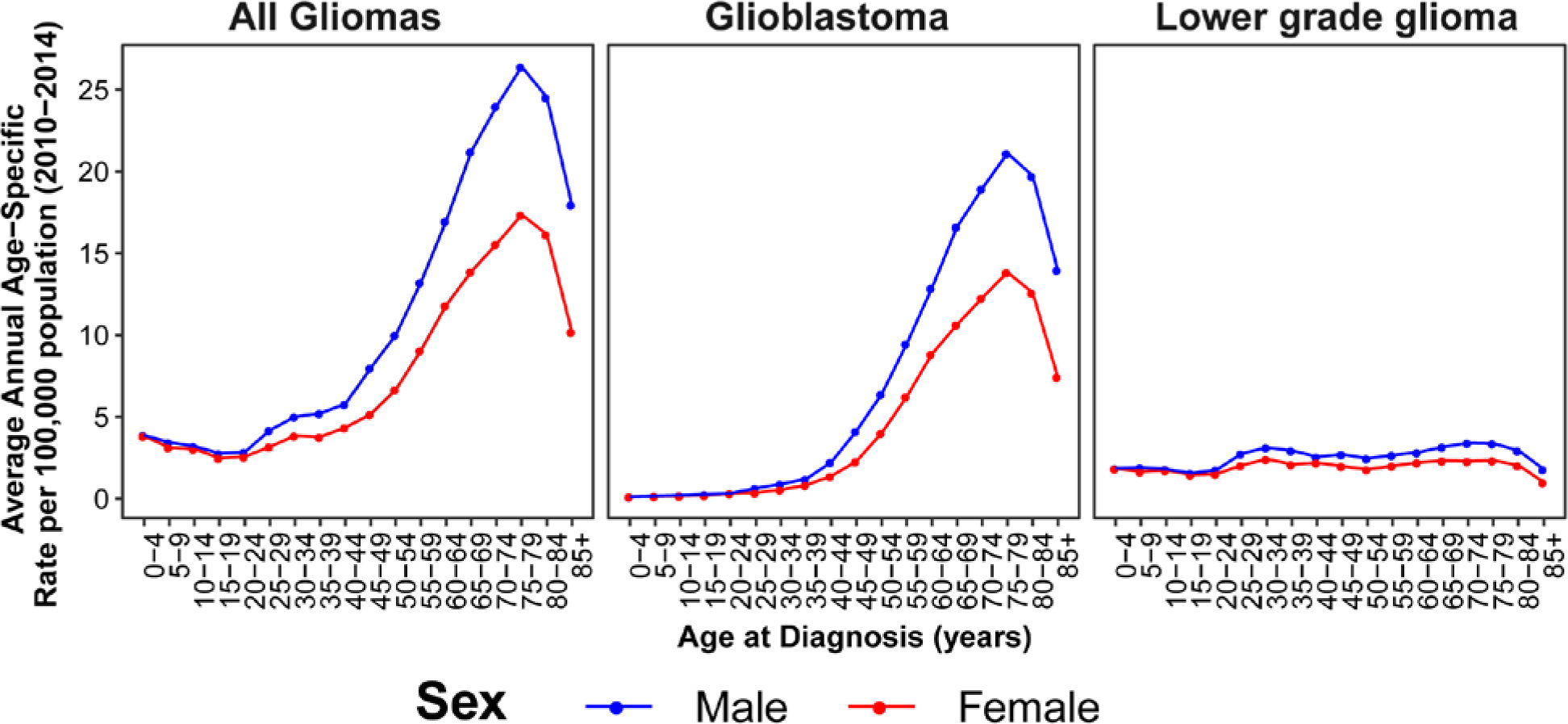
Average Annual Incidence of all glioma, glioblastoma and lower grade glioma by sex and age at diagnosis (CBTRUS 2010-2014)

Though sex differences exist in glioma incidence, sex differences have not been interrogated in previous glioma GWAS. Sex-specific analyses have the potential to reveal genetic sources of sexual dimorphism in risk, as well as to increase power for detection of loci where effect size or direction may vary by sex [9,10]. The aim of this analysis is to investigate potential sex-specific sources of genetic risk for glioma that may contribute to observed sex-specific incidence differences.

## Results

### Study population

There were 4,831 male cases, 5,216 male controls, 3,206 female cases, and 5,470 female controls (**Table 1**). A slightly larger proportion of male cases were GBM (58.7% of male cases vs 52.5% of female cases). Controls were slightly older than cases. GBM cases had a higher mean age than non-GBM cases, which was consistent with known incidence patterns of these tumors. Male and female cases within histology groups had similar age at diagnosis. The proportion of non-GBM cases varied by study due to differing recruitment patterns and study objectives (see original publications for details of recruitment patterns and inclusion criteria [4,11-14]).

**Table 1.** Population characteristics by study and sex

| Characteristic | Study | Males |  | Females |  |
| --- | --- | --- | --- | --- | --- |
|  |  | Cases | Controls | Cases | Controls |
| N | **Total** | **4,831** | **5,216** | **3,206** | **5,470** |
|  | GICC<sup>a</sup> | 2,733 | 1,868 | 1,831 | 1,397 |
|  | SFAGS-GWAS<sup>b</sup> | 440 | 749 | 237 | 1,611 |
|  | MDA-GWAS<sup>c</sup> | 714 | 1,094 | 429 | 1,142 |
|  | GliomaScan<sup>d</sup> | 944 | 1,465 | 709 | 1,260 |
| Mean Age (SD) | **Total** | **52.5 (14.5)** | **58.2 (15.2)** | **51.8 (14.9)** | **54.7 (14.5)** |
|  | GICC | 52.5 (14.3) | 56.1 (13.4) | 51.3 (14.6) | 53.4 (14.3) |
|  | SFAGS-GWAS | 53.8 (13.0) | 50.6 (14.8) | 53.5 (14.0) | 49.3 (13.2) |
|  | MDA-GWAS | 47.1 (13.0) | Modal age group: 60-69<sup>e</sup> | 47.7 (13.9) | Modal age group: 65-69<sup>f</sup> |
|  | GliomaScan | 56.0 (15.5) | 69.3 (12.7) | 55.1 (15.7) | 64.0 (15.4) |
| GBM (% of total)<sup>g</sup> | **Total** | **2,835 (58.7%)** | **--** | **1,682 (52.5%)** | **--** |
|  | GICC | 1,575 (57.6%) | -- | 885 (48.3%) | -- |
|  | SFAGS-GWAS | 333 (75.7%) | -- | 178 (75.1%) | -- |
|  | MDA-GWAS | 397 (55.6%) | -- | 246 (57.3%) | -- |
|  | GliomaScan | 530 (56.1%) | -- | 373 (52.6%) | -- |
| GBM - Mean Age (SD) | **Total** | **57.3 (12.0)** | **--** | **57.8 (12.1)** | **--** |
|  | GICC | 57.7 (11.4) | -- | 57.8 (11.6) | -- |
|  | SFAGS-GWAS | 56.4 (11.5) | -- | 56.2 (12.3) | -- |
|  | MDA-GWAS | 52.0 (11.7) | -- | 53.7 (11.3) | -- |
|  | GliomaScan | 60.4 (13.0) | -- | 61.4 (12.5) | -- |
| Non-GBM (% of total)<sup>g</sup> | **Total** | **1,716 (35.5%)** | **--** | **1,320 (41.2%)** | **--** |
|  | GICC | 1,036 (37.9%) | -- | 862 (47.1%) | -- |
|  | SFAGS-GWAS | 107 (24.3%) | -- | 59 (24.9%) | -- |
|  | MDA-GWAS | 317 (44.4%) | -- | 183 (42.7%) | -- |
|  | GliomaScan | 256 (27.1%) | -- | 216 (30.5%) | -- |
| Non-GBM - Mean Age (SD) | **Total** | **44.3 (14.4)** | **--** | **43.9 (14.3)** | **--** |
|  | GICC | 44.7 (14.6) | -- | 44.6 (14.2) | -- |
|  | SFAGS-GWAS | 45.7 (14.2) | -- | 45.4 (15.8) | -- |
|  | MDA-GWAS | 41.0 (11.9) | -- | 39.6 (12.9) | -- |
|  | GliomaScan | 46.3 (15.5) | -- | 44.4 (15.2) | -- |

a. Data from Glioma International Case-Control Study (GICC; Melin, et al.[4]); b. Data from San Francisco Adult Glioma Study GWAS (SFAGS-GWAS; Wrensch, et al.[12]); c. data from MD Anderson Cancer Center GWAS (MDA-GWAS; Shete, et al.[13]); d. Data from the National Cancer Institute's GliomaScan (GliomaScan; Rajaraman, et al.[14]); e. Data from CGEMS prostate study (Yeager et al. [15]). Continuous age is not available, age distribution is as follows 50-59: 12.3%, 60-69: 56.7%, 70-79: 30.7%, 80-89: 0.3%; f. Data from CGEMS breast study (Hunter et al. [16]). Continuous age is not available, age distribution is as follows: 0-54: 4.3%, 55-59: 15.0%, 60-64: 23.6%, 65-69: 27.5%, 70-74: 19.0%, 75-99: 10.7%; g. Histology information not available for all cases and frequencies may not add to 100%.

### Previously discovered glioma risk regions

There were 5,934 SNPs within 500kb of 26 previously discovered glioma risk loci with IMPUTE2 information score (INF0)>0.7 and MAF>0.01 that were previously found to have at least a nominal (p<5×10^−4^) association with glioma [4], and results were considered significant at p<2.8×10^−6^ level (adjusted for 6,000 tests in each of three histologies [18,000 tests], see **Figure 2a** for schematic of study design). Among the 25 previously validated glioma risk loci, nine loci contained 10 SNPs with p_M_<2.8×10^−6^ and/or p_F_<2.8×10^−6^ in any histology: 1p31.3 (*RAVER2*), 5p15.33 (*TERT*), 7p11.2 (EGFR, 2 independent loci), 8q24.21 (intergenic region near*MYC*), 9p21.3 (*CDKN2B-AS1*), 11q23.3 (*PHLDB1*), 16p13.3 (*RHBDF1*), 17p13.1 (*TP53*), and 20q13.33 (*RTEL1*) (**Table 2**). OR_M_ and OR_F_ were similar in the majority of these loci.

**Figure 2.**
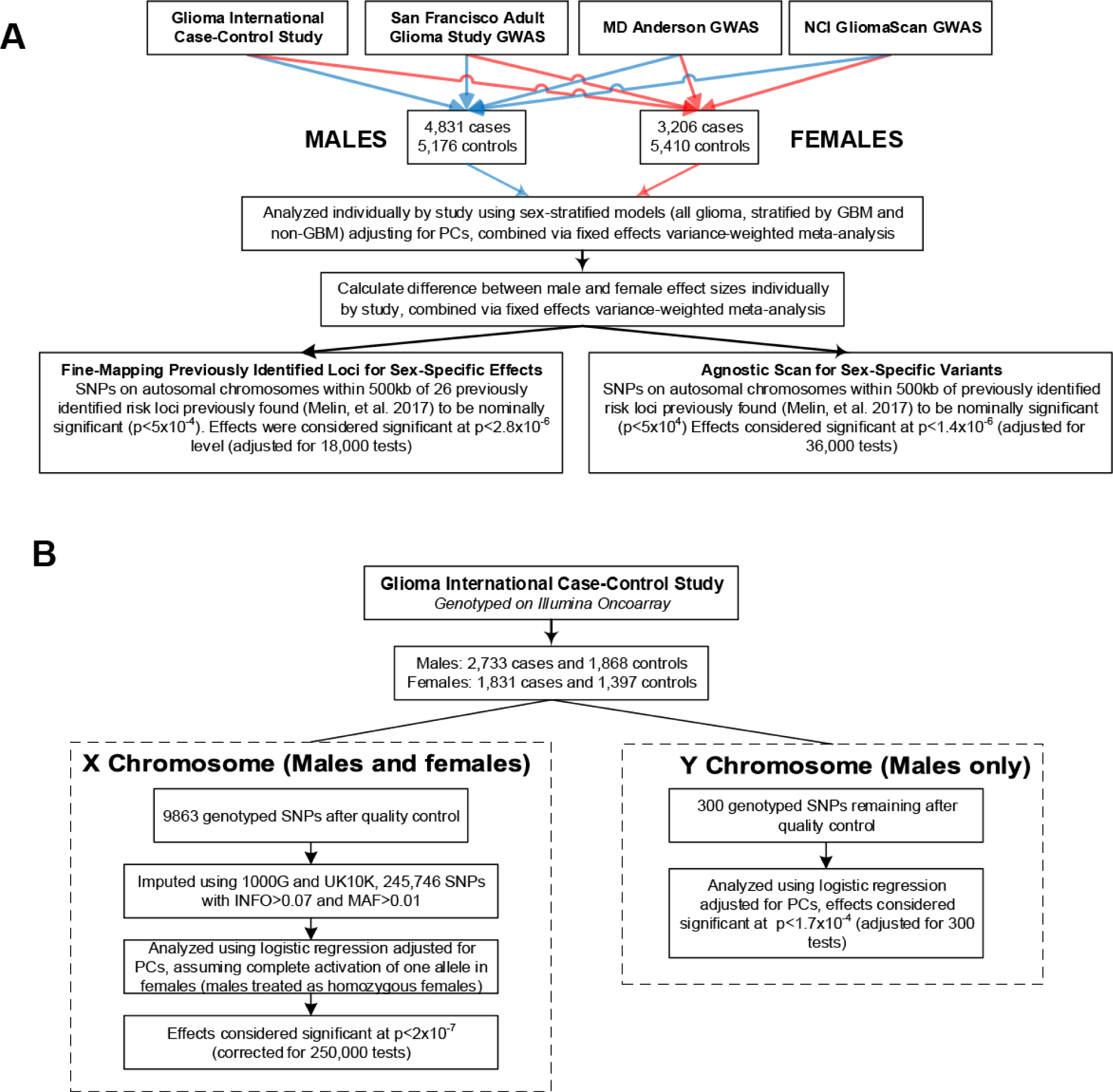
Study Schematic for analyses of a) autosomal SNPs and b) SNPs on sex chromosomes.

**Table 2.**
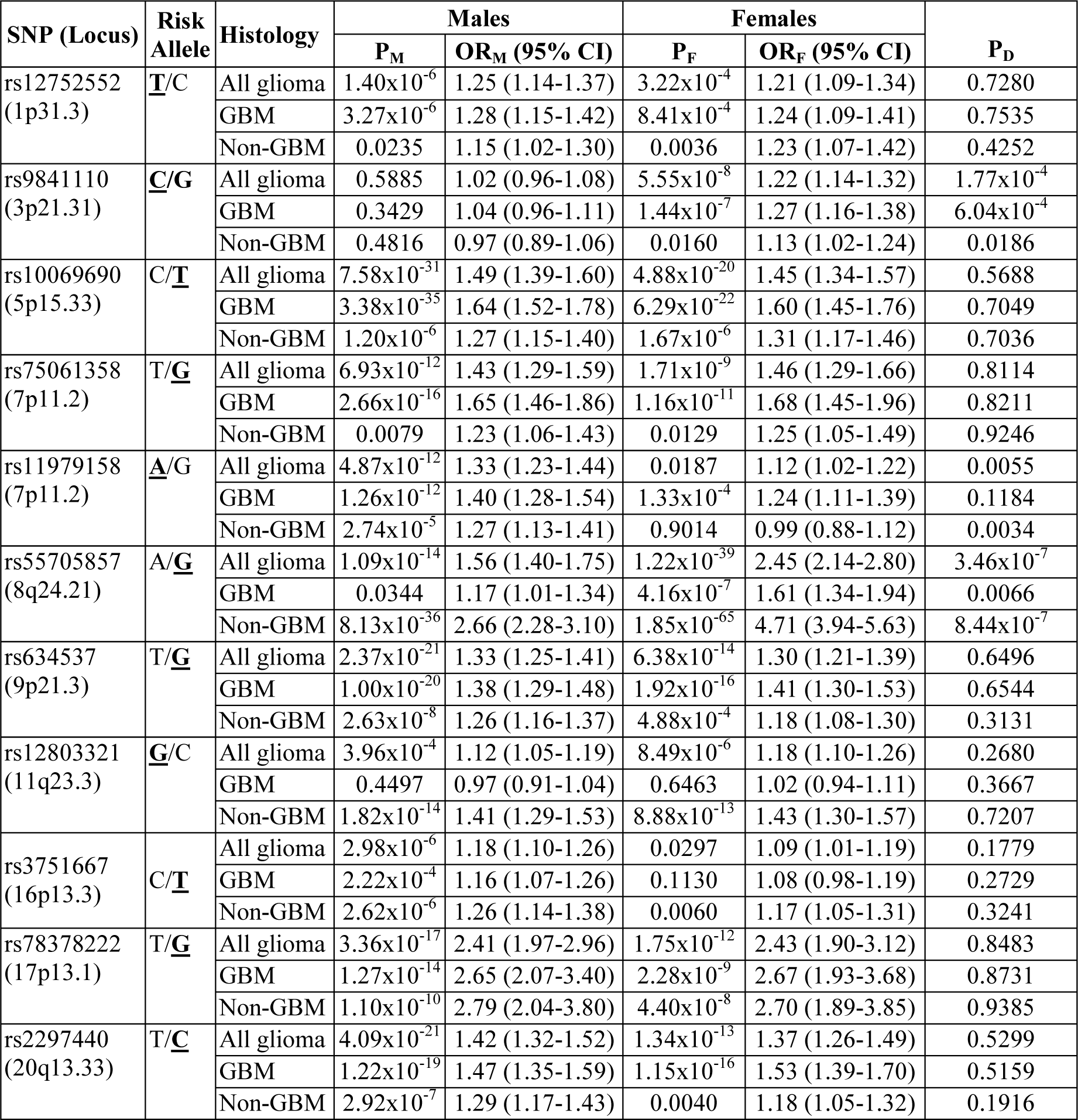
Previously identified glioma risk loci and histology-specific odds ratios (OR) and 95% confidence intervals (95% CI) stratified by sex.

For one of two independent loci at 7p11.2 (rs 11979158), there was a significant association only in males for all glioma (OR_M_= 1.33 [95% CI=1.23-1.44], p_M_=4.87×10^−12^) and GBM (OR_M_=1.40 [95% CI=1.28-1.54], p_M_=1.26×10^−12)^) but the sex differences did not meet the significance threshold (overall p_D_=0.0055, and GBM p_D_=0.1184) (**Figure 3, Table 2**).

**Figure 3.**
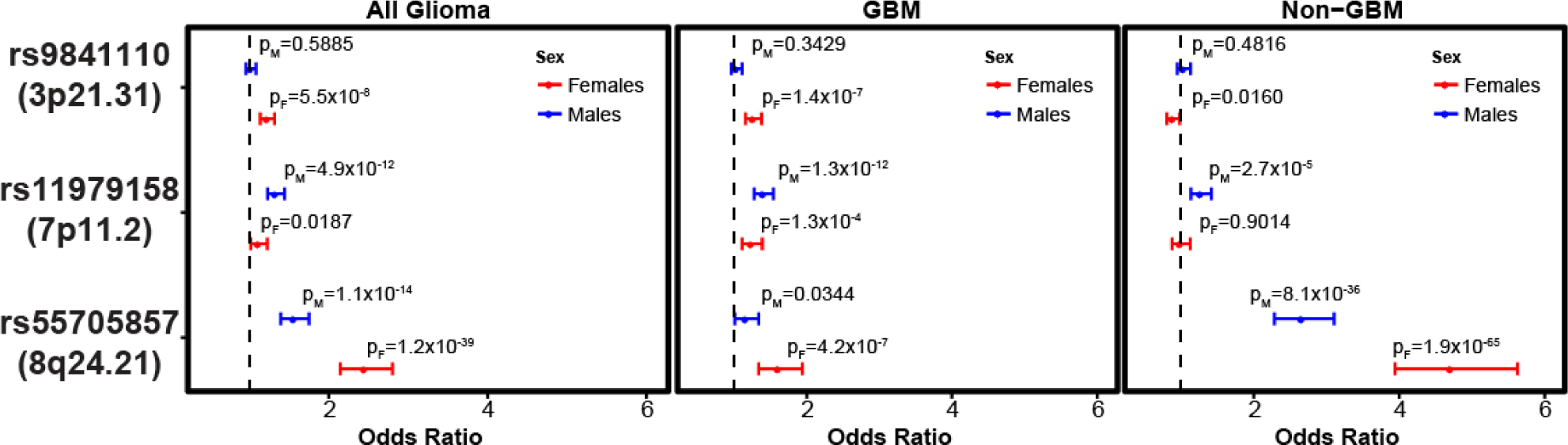
Sex-specific Odds ratios overall and by histology grouping, 95% CI and p values for selected previous GWAS hits and 3p21.31 (rs9841110) for all glioma, GBM, and non-GBM.

The previously identified SNP at 8q24.21 (rs55705857) was the most significant SNP in both males and females. Odds ratio for rs55705857 in all glioma was significantly higher in females (OR_F_=2.45 [95% CI=2.14-2.80], p_F_=1.22×10^−39^) as compared to males (OR_M_=1.56 [95% CI=1.40-1.75], pM=1.09×10^−14^) with p_D_=3.46×10^−7^. In non-GBM only, OR_F_ (OR_F_=4.71 [95% CI=3.94-5.63], p_F_=1.85×10^−65^) was also elevated as compared to OR_M_ (OR_M_=2.66 [95% CI=2.28-3.10)], p_M_=8.13×10^−36^) with p_D_=8.44x10^−7^ (**Figure 3, Table 2**). This association was further explored in a case-only analysis, where there was a significant difference between males and females overall (p=0.0012), and in non-GBM (p=0.0084) (**Table 3**).

**Table 3.**
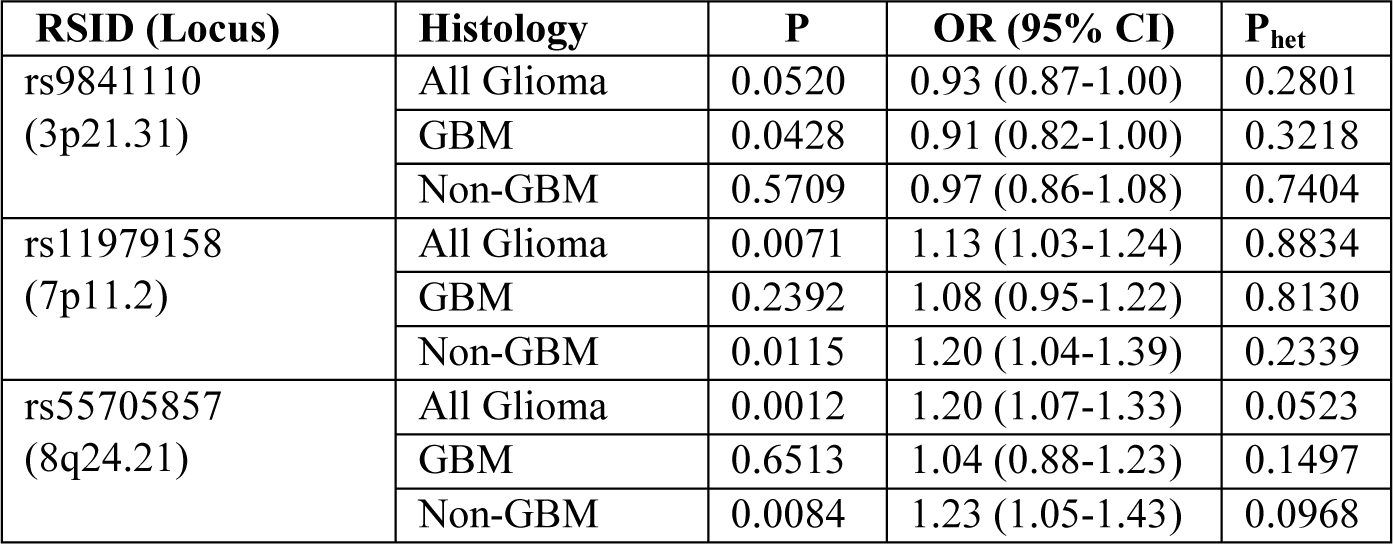
Case-only odds ratios (OR), 95% confidence intervals (95% CI), and p values from meta-analysis for rs11979158, rs55705857 and rs9841110 overall and by histology groupings.

| RSID (Locus) | Histology | P | OR (95% CI) | $P_{het}$ |
| --- | --- | --- | --- | --- |
| rs9841110<br>(3p21.31) | All Glioma | 0.0520 | 0.93 (0.87-1.00) | 0.2801 |
|  | GBM | 0.0428 | 0.91 (0.82-1.00) | 0.3218 |
|  | Non-GBM | 0.5709 | 0.97 (0.86-1.08) | 0.7404 |
| rs11979158<br>(7p11.2) | All Glioma | 0.0071 | 1.13 (1.03-1.24) | 0.8834 |
|  | GBM | 0.2392 | 1.08 (0.95-1.22) | 0.8130 |
|  | Non-GBM | 0.0115 | 1.20 (1.04-1.39) | 0.2339 |
| rs55705857<br>(8q24.21) | All Glioma | 0.0012 | 1.20 (1.07-1.33) | 0.0523 |
|  | GBM | 0.6513 | 1.04 (0.88-1.23) | 0.1497 |
|  | Non-GBM | 0.0084 | 1.23 (1.05-1.43) | 0.0968 |

Previous studies have found a strong association between rs55705857 and oligodendroglial tumors (particularly tumors with isocitrate dehydrogenase 1/2 (*IDH1/2*) mutation and loss of the 1p and 19q), so this association was further explored in the non-GBM (lower grade glioma [LGG]) histology groups (**Table 4**). For World Health Organization (WHO) grade II-grade III astrocytoma, effect was stronger in females (OR_F_=4.64 [95% CI=3.53-6.09], pF=2.15×10^−28^) as compared to males (OR_M_=2.87 [95% CI=2.31-3.56], p_M_=1.19×10^−21^) with p_D_=0.0065. For WH0 grade II-III Oligodendrogliomas effect was stronger than observed in WHO grade II-III astrocytomas, and effect size was stronger in females (OR_F_=12.15 [95% CI= 8.96-16.48], pF=3.68×10^−58^) as compared to males (OR_M_=5.47 [95% CI=4.16-7.19], pM=5.37×10^−34^) with p_D_=6.60×10^−5^. Oligoastrocytic tumors were not included in sub-analyses due to recent research that suggests that these tumors are not an entity that is molecularly distinct from oligodendrogliomas or astrocytomas [17].

**Table 4.** Sex-specific odds ratios (OR), 95% confidence intervals (95% CI), and p values from meta-analysis for rs11979158, rs55705857 and rs9841110 by specific non-GBM histologies.

| RSID<br>(Locus) | Histology | Males |  |  | Females |  |  | P<sub>D</sub> |
| --- | --- | --- | --- | --- | --- | --- | --- | --- |
|  |  | P<sub>M</sub> | OR<sub>M</sub><br>(95% CI) | P<sub>het</sub> | P<sub>F</sub> | OR<sub>F</sub><br>(95% CI) | P<sub>het</sub> |  |
| rs9841110<br>(3p21.31) | Astrocytoma (Non-GBM) (WHO grade II-III) | 0.5304 | 1.04 (0.92-1.17) | 0.751 | 0.0407 | 1.15 (1.01-1.32) | 0.549 | 0.2409 |
|  | Oligodendroglioma (WHO grade II-III) | 0.4190 | 0.94 (0.81-1.09) | 0.694 | 0.0973 | 1.14 (0.98-1.34) | 0.360 | 0.0649 |
| rs11979158<br>(7p11.2) | Astrocytoma (Non-GBM) (WHO grade II-III) | 0.0023 | 0.79 (0.68-0.92) | 0.056 | 0.9363 | 0.99 (0.83-1.18) | 0.418 | 0.0500 |
|  | Oligodendroglioma (WHO grade II-III) | 0.0221 | 0.81 (0.68-0.97) | 0.865 | 0.6561 | 1.05 (0.86-1.28) | 0.262 | 0.0471 |
| rs55705857<br>(8q24.21) | Astrocytoma (Non-GBM) (WHO grade II-III) | 1.19x10<sup>-21</sup> | 2.87 (2.31-3.56) | 0.073 | 2.15x10<sup>-28</sup> | 4.64 (3.53-6.09) | 0.237 | 0.0065 |
|  | Oligodendroglioma (WHO grade II-III) | 5.37x10<sup>-34</sup> | 5.47 (4.16-7.19) | 0.103 | 3.68x10<sup>-58</sup> | 12.15 (8.96-16.48) | 0.027 | 6.60x10<sup>-5</sup> |

### Genome-wide scan of nominally significant regions

In a previous eight study meta-analysis, ∼12,000 SNPs (INFO>0.7, MAF>0.01) were identified as having a nominally significant (p<5×10^−4^) association with all glioma, GBM, or non-GBM [4]. A sex-stratified genome-wide scan was conducted within this set of SNPs and results were considered significant at p_D_<1.4×10^−6^ (adjusted for 12,000 tests in each of three histologies [36,000 tests], see **Figure 2a** for schematic of study design). Similar genome-wide peaks were observed between males and females (**Figures 4-6**). One large region within 3p21.31 (49400kb-49600kb, ∼200kb) was identified as being significantly associated with glioma and GBM in females only (**Supplemental Figure 1**). There were 243 SNPs with nominally significant associations within this region in the previous eight-study meta-analysis (p<5×10^−4^), and 32 of these had nominally significant sex associations (p_F_<5×10^−6^ or p_M_<5×10^−6^) in all glioma or GBM. The strongest association in females within this region was at rs9841110, in both all glioma (OR_F_=1.22 [95% CI=1.14-1.32], pF=5.55×10^−8^) with pD=1.77×10^−4^) and GBM only (OR_F_=1.27 [95% CI=1.16-1.38], p_F_=3.86×10^−7^) with p_D_=6.04×10^−4^), while there were no significant associations detected in males (**Figure 3**). No SNPs in this region were significantly associated with non-GBM. In a case-only analysis a marginally significant difference was detected between males and females overall (p=0.0520) and in GBM (p=0.0428) (**Supplemental Table 1**).

**Figure 4.**
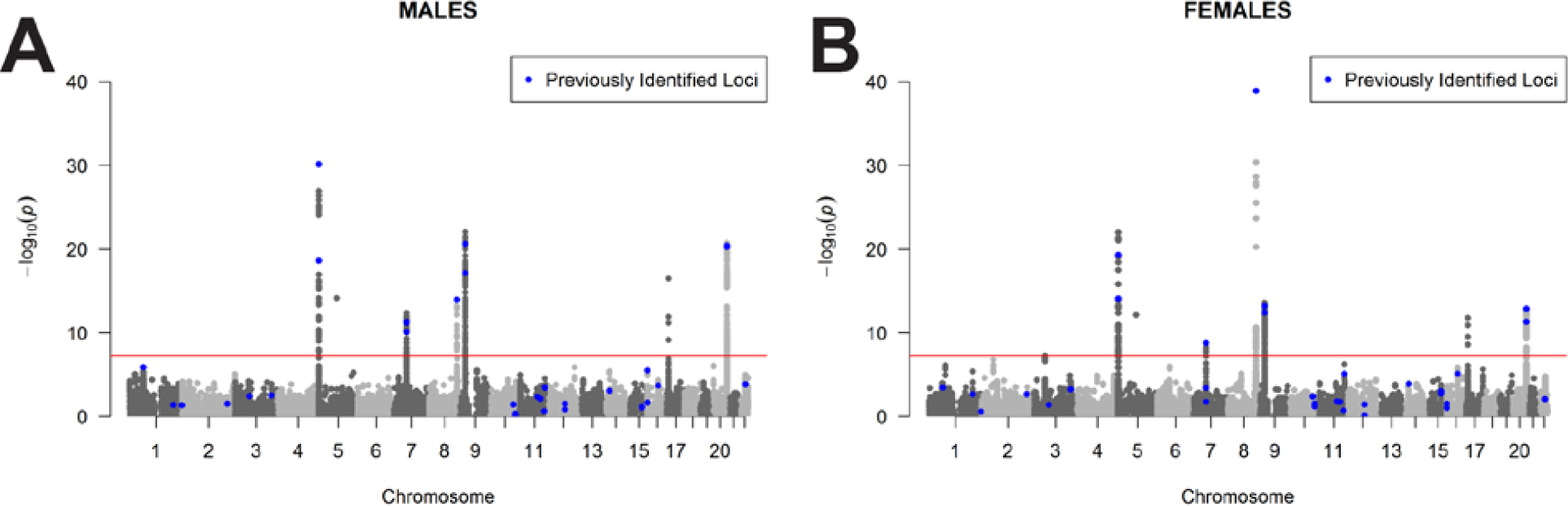
Manhattan plot of −log(p) values for all glioma in A) males and B) females.

**Figure 5.**
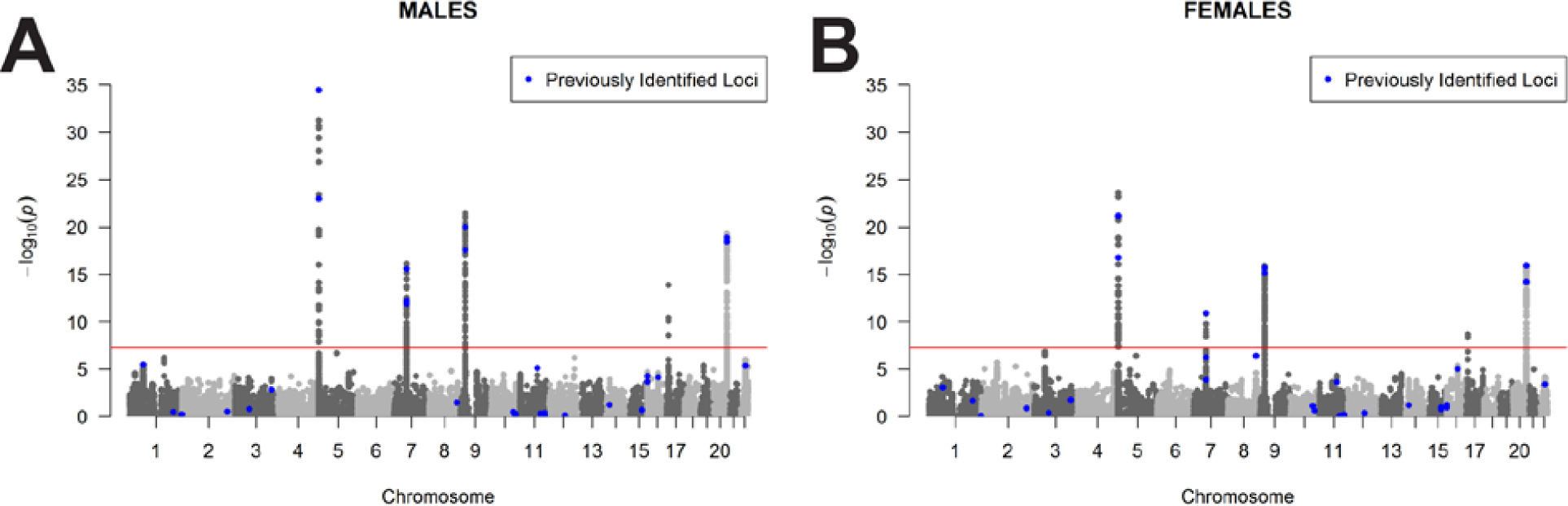
Manhattan plot of −log(p) values for GBM in A) males and B) females.

**Figure 6.**
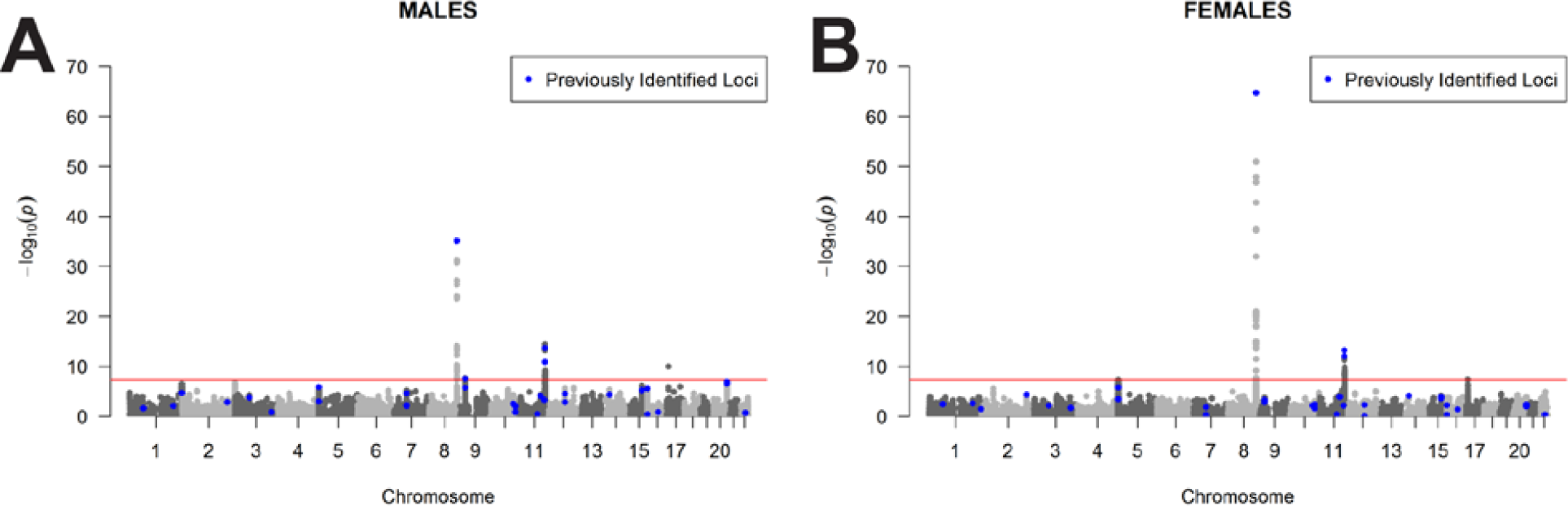
Manhattan plot of −log(p) values for non-GBM in A) males and B) females.

### Agnostic scan of sex chromosome loci

SNPs on the sex chromosomes were analyzed in GICC only. There were 245,746 SNPs with INFO>0.7 and MAF>0.01 on the X chromosome after quality control and imputation, and results were considered significant at p<2×10^−7^ (corrected for 250,000 tests, see **Figure 2b** for a schematic of study design). No SNPs met this significance threshold. After quality control procedures were complete, there were 300 SNPs remaining on the Y chromosome. No significant signals were detected on the Y chromosome.

### Combined analysis of germline variants and somatic characterization

Due to the lack of molecular classification data included in the GICC, MDA-GWAS, SFAGS-GWAS< and GliomaScan datasets, glioma data obtained from TCGA datasets (GBM and LGG) were used to explore the potential confounding due to molecular subtype variation with histologies. There were 758 individuals from the TCGA dataset available for analysis with available germline genotyping, molecular characterization, sex and age data (**Supplemental Table 2**). Overall, slightly more females (53.2%) as compared to males (47.2%) had *IDH1/2* mutant glioma, but this difference was not statistically significant (p=0.1104) (**Figure 7**). When tumors were stratified by histological type, approximately equal proportions of males and females had *IDH1/2* mutations present in their tumors (GBM: 6.0% in males, and 5.2% in females; LGG: 17.9% in males, and 17.7% in females). There were also no significant differences by sex in *IDH/TERT/*1p19q subtype (**Supplemental Figure 2**, overall p=0.2859), or panglioma methylation subgroup (**Supplemental Figure 3**, overall p=0.4153).

**Figure 7.**
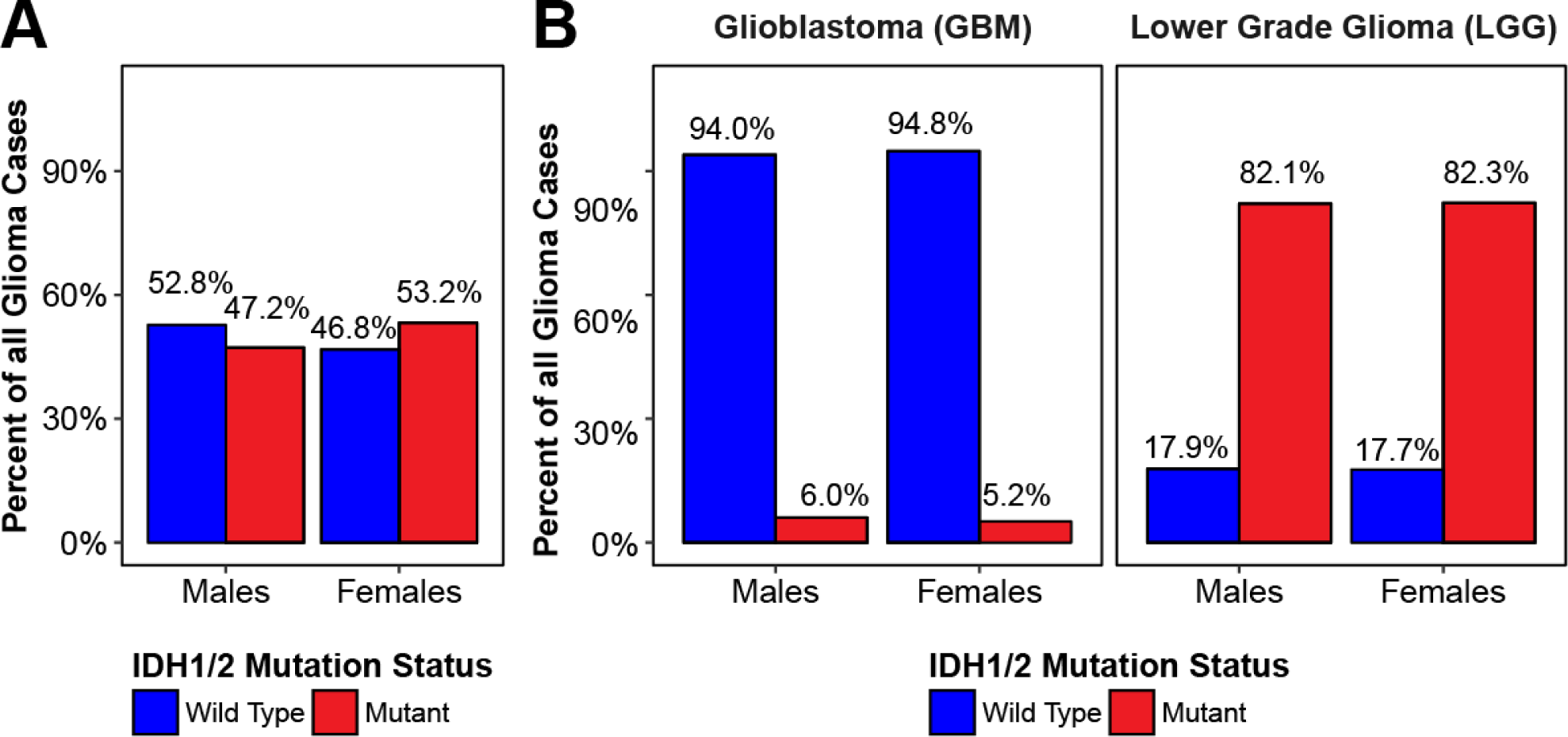
Proportion of samples with IDH1/2 mutation in the TCGA GBM and LGG datasets by sex, overall and stratified by study.

**Figure 8.**
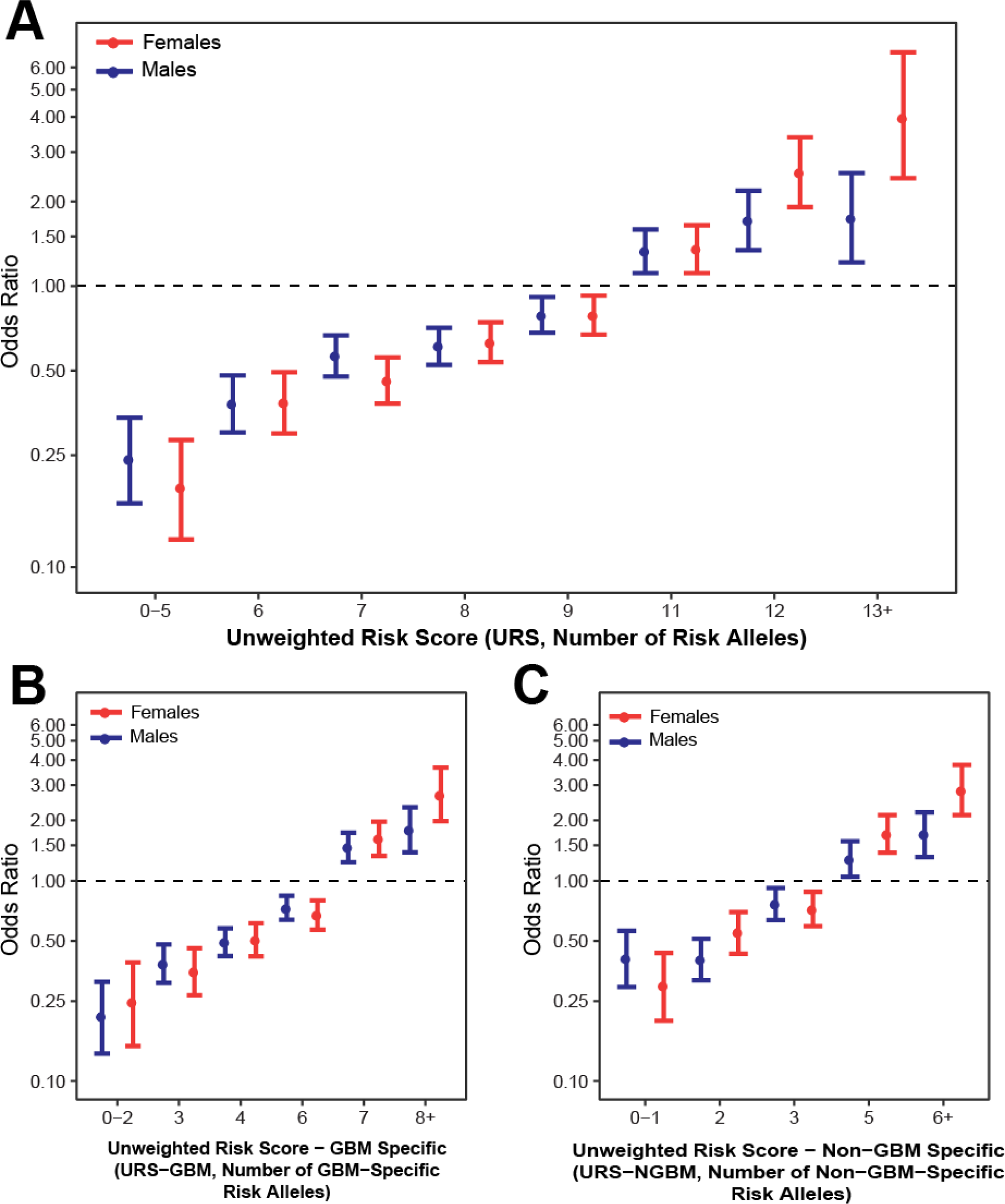
Odds ratios and 95% confidence intervals for unweighted risk (URS) score in A) all glioma, B)GBM-specific URS (URS-G) in GBM, and C) and non-GBM-specific URS (URS-NGBM) for in non-GBM.

SNPs found to be nominally significant (p<5×10^−4^) in a previous 8 study meta-analysis, with imputation quality (r^2^) ≥0.7 were identified within the TCGA germline genotype data and D’ and r^2^ values in CEU were used to select proxy SNPs (**Supplemental Table 3)** [18].

A case-only analysis was conducted using sex as a binary phenotype for proxy SNPs in the TCGA dataset. In the overall meta-analysis, there was a nominally significant signal in the case-only meta-analysis for the proxy SNP in 3p21.31 in glioblastoma (**Table 5**). There was no significant association in the TCGA set, but RAF was elevated in females as compared to males in the GBM set, as well as in all *IDH1/2* wild type gliomas (**Table 5**). MAF in LGG and *IDH1/2* mutant glioma was similar among males and females. There was a nominally significant signal in the case-only meta-analysis for the proxy SNP at 7p11.2, but no significant association in the TCGA, but RAF was elevated in males as compared to females in the GBM set, as well as in all *IDH1/2* wild type gliomas (**Table 5**). There was no significant signal detected in the overall case-only meta-analysis for the proxy SNP at 8q24.21, or within the TCGA set. Among both LGG and *IDH1/2* mutant, RAF was elevated in females as opposed to males.

**Table 5.** Risk allele frequencies (RAF) Case-only odds ratios, 95% confidence intervals (95% CI), and p values for marker SNPs from four study meta-analysis and the Cancer Genome Atlas genotyping data

| Marker SNP | Histology | Four-study Meta-Analysis |  |  |  | The Cancer Genome Atlas |  |  |  |  |
| --- | --- | --- | --- | --- | --- | --- | --- | --- | --- | --- |
|  |  | Males | Females | Case-only analysis (males:females) |  | INFO | Males | Females | Case-only analysis (males:females) |  |
|  |  | RAF<sub>cases</sub> | RAF<sub>cases</sub> | P | OR (95% CI) |  | RAF<sub>cases</sub> | RAF<sub>cases</sub> | p | OR (95% CI) |
| rs9814873<br>(3p21.31) | All glioma | 0.692 | 0.707 | 0.0577 | 1.07 (1.00-1.15) | 1.00 | 0.701 | 0.716 | 0.5321 | 0.93 (0.75-1.16) |
|  | GBM | 0.694 | 0.716 | 0.0371 | 1.11 (1.01-1.22) | 1.00 | 0.697 | 0.742 | 0.2003 | 0.80 (0.58-1.12) |
|  | LGG (non-GBM) | 0.686 | 0.691 | 0.6446 | 1.03 (0.92-1.15) | 1.00 | 0.705 | 0.697 | 0.8039 | 1.04 (0.77-1.40) |
|  | *IDH1/2* wild type | -- | -- | -- | -- | 1.00 | 0.704 | 0.731 | 0.4343 | 0.88 (0.64-1.21) |
|  | *IDH1/2* mutant | -- | -- | -- | -- | 1.00 | 0.705 | 0.692 | 0.7023 | 1.06 (0.77-1.47) |
| rs7785013<br>(7p11.2) | All glioma | 0.864 | 0.847 | 0.0058 | 0.88 (0.80-0.96) | 0.99 | 0.855 | 0.850 | 0.7813 | 1.04 (0.78-1.39) |
|  | GBM | 0.872 | 0.861 | 0.2141 | 0.92 (0.81-1.05) | 0.99 | 0.854 | 0.840 | 0.6073 | 1.12 (0.73-1.72) |
|  | LGG (non-GBM) | 0.855 | 0.832 | 0.0109 | 0.83 (0.72-0.96) | 0.99 | 0.856 | 0.857 | 0.9585 | 0.99 (0.67-1.47) |
|  | *IDH1/2* wild type | -- | -- | -- | -- | 0.99 | 0.864 | 0.837 | 0.3132 | 1.24 (0.82-1.87) |
|  | *IDH1/2* mutant | -- | -- | -- | -- | 0.99 | 0.846 | 0.875 | 0.2447 | 0.77 (0.50-1.19) |
| rs4636162<br>(8q24.21) | All glioma | 0.358 | 0.365 | 0.5161 | 1.02 (0.96-1.09) | 0.93 | 0.392 | 0.424 | 0.2113 | 0.88 (0.71-1.08) |
|  | GBM | 0.343 | 0.338 | 0.6001 | 0.98 (0.89-1.07) | 0.93 | 0.374 | 0.404 | 0.4456 | 0.89 (0.66-1.20) |
|  | LGG | 0.383 | 0.401 | 0.1594 | 1.08 (0.97-1.20) | 0.94 | 0.410 | 0.438 | 0.3891 | 0.88 (0.67-1.17) |
|  | IDH1/2<br>wild type | -- | -- | -- | -- | 0.92 | 0.371 | 0.373 | 0.9480 | 0.99 (0.73-1.34) |
|  | IDH1/2<br>mutant | -- | -- | -- | -- | 0.94 | 0.419 | 0.460 | 0.2613 | 0.84 (0.63-1.14) |

### Sex-stratified genotypic risk scores

In order to estimate the cumulative effects of significant variants by sex, unweighted risk scores (URS) were calculated by summing all risk alleles for each individual using the 10 SNPs (rs12752552, rs9841110, rs10069690, rs11979158, rs55705857, rs634537, rs12803321, rs3751667, rs78378222, and rs2297440) found to be significantly associated with glioma in this analysis. GBM (URS-GBM) and non-GBM (URS-NGBM) specific URS were calculated only using sets of 6 SNPs in this set that were significantly associated with these histologies (URS-GBM: rs9841110, rs10069690, rs11979158, rs634537, rs78378222, and rs2297440, and URS-NGBM: rs10069690, rs55705857, rs634537, rs12803321, rs78378222, and rs2297440). See Methods for additional information on score calculation. Median URS, URS-GBM, and URS-NGBM were significantly different (p<0.0001) between cases and controls in both males and females in all histology groups (**Supplemental Figure 4**). There was no significant difference in median risk scores between male and female cases for any histology group. Glioma risk increased with increasing number of alleles in both males and females for the 10 SNPs included in the overall URS, as well as the 6 SNPs in the URS-GBM and 6 SNPs in URS-NGBM (**Figure 9**, **Supplemental Table 4**). Risk was higher in females (OR=3.97 [95% CI=2.42-6.80]) as compared to males (OR=1.74 [95% CI=1.21-2.53]) in all glioma for individuals for with 13-16 alleles, though the difference between these estimates were not statistically significant. Risk was also higher among females (OR=2.69 [95% CI=1.98-3.66]) as compared to males (OR=1.79 [95% CI=1.38-2.32]) in GBM for individuals with 8-11 risk alleles, as well as in non-GBM for individuals with 6-11 risk alleles (females: OR=2.83 [95% CI=2.12-3.78], males: OR=1.70 [95% CI=1.31-2.19]), though the difference between these estimates were not statistically significant. The estimates may underestimate actual risk due to varying effect sizes and alleles frequencies between risk variants.

## Discussion

This is the first analysis of inherited risk variants in sporadic glioma focused specifically on sex differences, and the first agnostic unbiased scan for glioma risk variants on the X and Y sex chromosomes. Like many other types of chronic disease, there is a male preponderance of glioma. This incidence difference is not currently explained by known environmental or genetic risk factors.

One SNP at the 7p11.2 locus (rs11979158) showed significant association in males only, in both all glioma and GBM (**Table 2**). Effects were similar in all studies included in the analysis (**Supplemental Table 5**, **Supplemental Figure 6**). This variant is within one of two previously identified independent glioma risk loci located near epidermal growth factor receptor (*EGFR*) and is most strongly associated with risk for GBM.[4,19] Though *EGFR* is implicated in many cancer types and is a target for many anti-cancer therapies, this risk locus has not been previously associated with any other cancer type. Estrogen has been demonstrated to interact with *EGFR* as well as other growth factors [20]. Previous studies have not been definitive about the role of endogenous estrogen exposure in glioma risk, so it was not possible to determine the biological plausibility of this association [20]. Alternatively, cell intrinsic, hormone independent sex differences in EGF effects have been observed in a murine model of gliomagenesis in which EGF treatment was transforming for male but not female astrocytes that had been rendered null for neurofibromin and p53 function [21]. While this specific SNP was not genotyped on the germline genotyping array used for TCGA, a SNP in strong LD with rs11979158 (rs7785013, D’=1, r^2^=1 in CEU [18]) was evaluated. The association in the case-only analysis in TCGA was not statistically significant in any histology group, but a similar trend to that observed in the overall meta-analysis in sex-specific RAF was observed in both the overall GBM group, as well as in the *IDH1/2* wild type group.

The association at 8q24.21 (rs55705857) is the strongest that has been identified by glioma GWAS to date,[4] with an odds ratio of 1.99 (95% CI=1.85-2.13, p=9.53×10^−79^) in glioma overall, and an odds ratio of 3.39 (95% CI=3.09-3.71, p=7.28×10^−149^) in non-GBM. Effects were similar in all studies included in the analysis (**Supplemental Table 5**, **Supplemental Figure 7**). The identified SNP, rs55705857, is located in an intergenic region near coiled-coil domain containing 26 (*CCDC26*, a long non-coding RNA). This analysis found a stronger association in females than males in all glioma and non-GBM, where female odds ratio estimates are ∼2x those of males (Table 2). ORs were higher in women than men in all studies included in the analysis, but the magnitude of the ORs varied between studies (**Supplemental Table 5**). Furthermore, the MAF for rs55705857 in the SFAGS-GWAS differed from the other three studies (See **Supplemental Table 6** for MAF by study). Consequently, a sensitivity analysis was conducted to assess the effect of study heterogeneity on this estimate in non-GBM using only the GICC, MDA-GWAS, and GliomaScan datasets. The results of this analysis did not substantially change from (Main analysis p_D_=1.20×10^−6^ and sensitivity p_D_=1.49×10^−5^).

A histology-specific analysis found a similar sex differences in ORs for rs55705957 for both non-GBM astrocytoma, and oligodendroglioma (**Table 4**, see **Supplemental Table 7** for study-specific estimates). Previous analyses have shown that this variant is strongly associated with *IDH1/2* mutant tumors, particularly those that have 1p/19q deletions [22,23]. Data on *IDH1/2* mutation and 1p/19q codeletion were not available for the combined four GWAS datasets used here. Hence, to assess potential differences in frequency of *IDH1/2* mutation, the frequency of these mutations by sex was assessed within the combined TCGA GBM and LGG datasets [24-26]. Approximately the same proportion of males as females with histologically confirmed GBM had *IDH1/2* mutations (5.2% vs 6.0%, respectively), so females may not be more likely than males to present with *IDH1/2* mutant GBM (**Figure 7**). While this specific SNP was not genotyped on the germline genotyping array used for TCGA, a SNP in weak LD with rs55705857 (rs4636162, D’=1; r^2^=0.104, in CEU [18]) was able to be evaluated. There was no significant association in the overall case-only meta-analysis for this SNP, and the association in the case-only analysis in TCGA was not statistically significant in any histology group. Sex-specific RAF for this SNP was slightly higher in females as compared to males in the overall LGG group as well as the *IDH1/2* mutant group.

A large region in 3p21.31 was identified that was associated with all glioma and GBM in females only (Table 2). Effects were similar in all studies included in the analysis (**Supplemental Table 5**, **Supplemental Figure 7**). The strongest association in this region was rs9841110, an intronic variant located upstream of dystroglycan 1 (*DAG1*) within an enhancer region. While this specific SNP was not genotyped on the germline genotyping array used for TCGA, a SNP in strong LD with rs9841110 (rs9814873, D’=1, r^2^=1 in CEU [18]) was able to be evaluated. The association in the case-only analysis in TCGA was not statistically significant in any histology group, but a similar trend to that observed in the overall meta-analysis in sex-specific RAF was observed in both the overall GBM group, as well as in the *IDH1/2* wild type group. The identified risk allele at rs9841110 (C) is associated with significantly increased expression in glutathione peroxidase 1 (*GPX1* 1.3×10^−7^ in cerebellum, p=3.1×10^−7^, in frontal cortex), macrophage stimulating 1 receptor (*MST1R* [RON], p=4.3×10^−5^ in cerebellar hemisphere) and ring finger protein 123 (*RNF123* [*KPC1*), and significantly decreased expression of macrophage stimulating 1 (*MST1*, p=4.8×10^−6^ in hypothalamus and –p=1.5×10^−5^ in cerebellum),and RNA binding motif protein 6 (*RBM6*, p=2.7×10^−6^ in cerebellum) in normal brain tissue [27]. Glioblastoma samples have elevated expression of *GPX1* (fold change 2.79) and decreased expression of *MST1R* (fold change 0.44) as compared to normal tissue[25,28], and increased expression of GPX1 and MSTIR1 have been associated with poor prognosis in multiple cancer types [29,30].

Though this region has not previously been associated with glioma, previous GWAS have detected associations at 3p21.31 for a large variety of traits, including several autoimmune diseases as well as increased age at menarche [31-34]. Three variants previously associated with increased age at menarche (rs7647973-A: D’=1.0 and r^2^=0.1441; rs6762477-G: D’=0.66 and r^2^=0.1659; rs7617480-A: D’=1.0 and r^2^=0.1332) are in linkage disequilibrium with the identified risk allele at rs9841110 (C), with in CEU [18,32]. If lifetime estrogen exposure modifies glioma risk, it is reasonable that variants which increase age at menarche, which may potentially decrease total lifetime estrogen exposure, may also be related to glioma risk in females. Due to the complexity of measuring lifetime estrogen exposure (which is affected by age at menarche, age at menopause, parity, breast feeding patterns, and estrogen replacement therapy post-menopause) it is difficult to determine the ‘true’ effect that this exposure might have on glioma risk.

As compared to a model containing age at diagnosis and sex alone, the three SNPs (rs55705857, rs9841110 and rs11979158) identified as having sex-specific effects explain an additional 1.4% of trait variance within the GICC set. The variance explained by these SNPs varies by histology (0.6% in GBM, and 3.3% in Non-GBM). The variance explained by the addition of these three SNPs was higher in females for all glioma (1.3% in males and 2.2% in females), and non-GBM glioma (2.3% in males and 5.3% in females), and slightly higher in males for GBM (0.9% in males and 0.7% in females).

In order to compare the cumulative effects of glioma risk variants by sex, unweighted risk scores (URS) were generated by summing all risk alleles using the 10 SNPs found to be significantly associated with glioma in this analysis. GBM (URS-GBM) and non-GBM (URS-NGBM) specific URS were calculated using sets of 6 SNPs in this set that were associated with significantly associated with these histologies. Individuals with lower numbers of risk alleles had significantly lower risk of glioma, and those with higher numbers of alleles had increased risk for glioma, with statistically significant trends in each histology group). Males and females with low risk scores had similar odds of glioma, while females had increased odds in the upper strata of scores as compared to males. Development of risk scores that weight alleles by effect size, and use sex-specific estimates for variants for which effect size varies by sex (such as 7p11.2 and 8q24.21), may lead to better predictive values for risk scores.

This is the first sex-specific analysis of germline risk variants for glioma, and identifies three loci with sex-specific effects, and leverages multiple existing glioma GWAS datasets. While often not included in GWAS, sex-stratified analyses can reveal genetic sources of sexual dimorphism in risk, [9,10]. Sex variation in genetic susceptibility to disease is likely not due to sex differences in DNA sequence, but is likely to be related to sex-specific regulatory functions [35-37]. These analyses may not only contribute to understanding of sources of sex difference in incidence, but may also suggest mechanisms and pathways that vary by sex in contributions to gliomagenesis.

In addition to genetic sources of difference, there are likely several additional factors acting in combination which contribute to sex differences in glioma incidence. Sex differences in disease can also be linked to in-*utero* development, during which time gene expression and risk phenotypes are patterned through the action of X alleles that escape inactivation and genes on the non-pseudo-autosomal component of the Y chromosome, as well as the epigenetic effects of in utero testosterone. [38]. A previous analysis estimating heritability of brain and CNS tumors by sex using twins attempted to estimate sex-specific relative risks, but these analyses were limited by a small sample size [39]. Further investigation of the inheritance patterns of familial glioma by sex may also provide additional information about sex differences in this disease.

There are several limitations to this analysis. Individuals included in these datasets were recruited during different time periods from numerous institutions, with no central review of pathology. Molecular tumor markers were unavailable for all datasets, and as a result classifications are based on the treating pathologist using the prevailing histologic criteria at time of diagnosis. The variant at 8q24.21 has been shown to have significant association with particular molecular subtypes, and without molecular data it was not possible to determine whether the observed result is an artifact of varying molecular features by sex. Oligodendroglioma as a histology is highly enriched for *IDH1/2* and 1p/19q co-deleted tumors (117/174, or ∼67% within the TCGA glioma dataset [24] and it is therefore likely that the analysis using only tumors classified as oligodendroglioma captured most of this molecular subtype. Males and females within histology groups have different frequencies of *IDH1/2* mutation [24], which may have confounded the estimates for 8q24.21. The TCGA dataset was used to explore sex differences in allele frequency within molecular groups, but none of the identified SNPs were able to be directly validated within this set; however SNPs in strong LD were evaluated except for in 8q24.21. The 8q24.21 region is not well characterized on the array used for the TCGA genotyping, and as a result this region imputed poorly. No proxy SNP in strong LD with rs55705857 was able to be identified. Similar trends in RAF to those observed in the overall meta-analysis were seen in the TCGA set, though these differences were not statistically significant. Further interrogation in datasets with molecular classification where direct genotyping of these regions is warranted in order to confirm the sex-specific associations observed in this analysis.

## Conclusions

Sex and other demographic differences in cancer susceptibility can provide important clues to etiology, and these differences can be leveraged for discovery in genetic association studies. This analysis identified potential sex-specific effects in 2 previous identified glioma risk loci (7p11.2, and 8q24.21), and 1 newly identified autosomal locus (3p21.31). Odds ratios for the highest strata of an unweighted risk score calculated by summing total risk alleles was higher in females as compared to males in all three histology groups. These significant differences in effect size may be a result of differing biological function of these variants by sex due to biological sex differences, or interaction between these variants and unidentified risk factors that vary in prevalence or effect by sex.

## Materials and Methods

### Study cohorts

This study was approved locally by the institutional review board (IRB) at University Hospitals Cleveland Medical Center and by each participating study site’s IRB. Written informed consent was obtained from all participants. In this study, data was combined from four prior glioma GWAS: Glioma International Case-Control Study (GICC), San Francisco Adult Glioma Study GWAS (SFAGS-GWAS), MD Anderson Glioma GWAS (MDA-GWAS), and National Cancer Institute’s GliomaScan (**Figure 4A**) [4,11-14]. The SFAGS-GWAS includes controls from the Illumina iControls dataset, and MDA-GWAS includes controls from Cancer Genetic Markers of Susceptibility (CGEMS) breast and prostate studies [15,16,40]. Details of data collection and classification are available in previous publications [4,11-14].

### Genotyping and imputation of GWAS datasets

GICC cases and controls were genotyped on the Illumina Oncoarray [41]. The array included 37,000 beadchips customized to include previously-identified glioma-specific candidate single nucleotide polymorphisms (SNPs). SFAGS-GWAS cases and some controls were genotyped on Illumina’s HumanCNV370-Duo BeadChip, and the remaining controls were genotyped on the Illumina HumanHap300 and HumanHap550. MDA-GWAS cases were genotyped on the Illumina HumanHap610 and controls using the Illumina HumanHap550 (CGEMS breast [16,40]) or HumanHap300 (CGEMS prostate [15]). GliomaScan cases were genotyped on the Illumina 660W, while controls were selected from cohort studies and were genotyped on Illumina 370D, 550K, 610Q, or 660W (See Rajaraman et al. for specific details of genotyping) [14]. Details of DNA collection and processing are available in previous publications [4,12-14]. Individuals with a call rate (CR) <99% were excluded, as well as all individuals who were of non-European ancestry (<80% estimated European ancestry using the FastPop [42] procedure developed by the GAMEON consortium)). For all apparent first-degree relative pairs were removed (identified using estimated identity by descent [IBD]≥.5), for example, the control was removed from a case-control pair; otherwise, the individual with the lower call rate was excluded. SNPs with a call rate <95% were excluded as were those with a minor allele frequency (MAF)<0.01, or displaying significant deviation from Hardy-Weinberg equilibrium (HWE) (p<1×10^−5^). Additional details of quality control procedures have been previously described in Melin et al [4]. All datasets were imputed separately using SHAPEIT and IMPUTE using a merged reference panel consisting of data from the 1,000 genomes project and the UK10K [43-47].

TCGA cases were genotyped on the Affymetrix Genomewide 6.0 array using DNA extracted from whole blood (see previous manuscript for details of DNA processing [25,26]), and underwent standard GWAS QC, and duplicate and related individuals within datasets have been excluded [4]. Ancestry outliers were identified in TCGA using principal components analysis in plink 1.9 [48]. Resulting files were imputed using Eagle 2 and Minimac3 as implemented on the Michigan imputation server (https://imputationserver.sph.umich.edu) using the Haplotype Reference Consortium Version r1.1 2016 as a reference panel [49-51]. Somatic characterization of TCGA cases was obtained from the final dataset used for the TCGA pan-glioma analysis [24], and classification schemes were adopted from Eckel-Passow, et al.[52] and Ceccarelli, et al.[24].

### Sex-stratified scan of the autosomal chromosomes

The data were analyzed using sex-stratified logistic regression models in SNPTEST for all SNPs on autosomal chromosomes within 500kb of previously identified risk loci, and/or those found to be nominally significant (p<5×10^−4^) in a previous meta-analysis (**Figure 2A**) [4,53]. Sex-specific betas (β_M_ and β_F_), standard errors (SE_M_ and SE_F_), and p-values (p_M_ and p_F_) were generated using sex-stratified logistic regression models that were adjusted for number of principal components that significantly differed between cases and controls within each study.

### Estimation of sex difference and test of statistical significance

β_D_ and SE_D_ were estimated using the sex-specific betas and standard errors separately for each dataset, as follows:

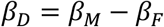

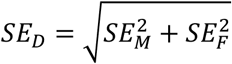

The difference between the groups was then tested using a z test.[54,55] Sex-stratified results and differences estimates from the four studies were separately combined via inverse-variance weighted fixed effects meta-analysis in META [56]. See **Figure 2A** for schematic of autosomal analysis methods. Case only-analyses were performed for SNPs found to be significant in agnostic analyses using sex as outcome for all glioma, GBM, and non-GBM by study and betas and standard errors were combined via inverse-variance weighted fixed effects meta-analysis in META [56].

### Sex chromosome analysis

X and Y chromosome data were available from GICC set only. Males and females were imputed separately for the X chromosome using the previously described merged reference panel. X chromosomes were analyzed using logistic regression model in SNPTEST module ‘newml’ assuming complete inactivation of one allele in females, and males are treated as homozygous females (**Figure 2B**). For prioritized SNPs in the combined model, sex-specific effect estimates were generated using stratified logistic regression models. Y chromosome data were analyzed using logistic regression in SNPTEST (**Figure 2B**) [57]. Figures were generated using R 3.3.2, GenABEL, qqman, and ggplot.[58-61]

### Analysis of TCGA germline and somatic data

Only newly diagnosed cases from TCGA GBM and LGG with no neo-adjuvant treatment or prior cancer were used. Demographic characteristics, molecular classification and somatic alterations data was obtained from Ceccarelli, et al [24]. Chi-square tests were used to compare the frequency of somatic alterations between age groups. SNPs found to be nominally significant (p<5×10^−4^) in a previous 8 study meta-analysis [4], with imputation quality >= 0.7 were identified within the TCGA genotype data and D’ and r^2^ values in CEU were used to select proxy SNPs [18]. Using these SNPs, a case-only analysis using sex as a binary phenotype was conducted using logistic regression in SNPTEST assuming an additive model to estimate beta, standard error, and p values [53]. Results were considered significant at p<0.003 (Bonferroni correction for 15 tests, for the three assessed loci in each of five histology groups).

### Calculation of unweighted genetic risk scores

In order to estimate the cumulative effects of significant variants by sex, histology-specific unweighted risk scores were calculated using the SNPs found to be significantly associated with each outcome. Data from all four studies was merged, and any imputed genotypes with genotype probability > 0.8 were converted to hard calls. An overall unweighted risk score (URS) was generated using the sum of risk alleles at rs12752552, rs9841110, rs10069690, rs11979158, rs55705857, rs634537, rs12803321, rs3751667, rs78378222, and rs2297440. As risk alleles are known to have histology specific associations,[4] histologic specific scores were generated for GBM and non-GBM using only the SNPs found to have a significant association with each histology. GBM-specific URS (URS-G) was calculated by summing the number of risk alleles at rs9841110, rs10069690, rs 11979158, rs634537, rs78378222, and rs2297440. Non-GBM-specific (URS-N) specific URS was calculated by summing the number of risk alleles at rs10069690, rs55705857, rs634537, rs12803321, rs78378222, and rs2297440. Unweighted risk scores (URS) were calculated by summing all risk alleles for each individual. Differences in median scores between groups using were tested using Wilcoxon rank sum tests. Scores were compared against the median score for each set (URS: 10, URS-GBM: 6 alleles, URS-NGBM: 4 alleles). Odds ratios and 95% confidence intervals for each level of the score using sex-stratified logistic regression adjusted for age at diagnosis (for controls where only an age range was available, the mean value of the range was used), where each score was compared to the median score within the entire population as described in Shete et al. [13].

### Calculation of trait variance explained by SNPs with sex-specific effects

In order to determine whether the identified SNPs with sex-specific effects more accurate estimate odds of glioma than sex alone, logistic regression models were used to estimate odds of all glioma, GBM, and non-GBM glioma based on sex using the GICC data only. Proportion of variance in odds of glioma explained by sex-specific SNPs was calculated using R^2^ estimated using the log likelihood of the null model (sex, age at diagnosis, and the first two principal components only) and the full model (including identified SNPs, rs9841110, rs11979158, rs55705857) [62], calculated as follows:

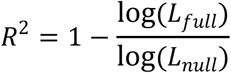

Proportion of variance explained was also calculated separately by sex for each histology (null model adjusted for age at diagnosis, and the first two principal components only).

## Acknowledgements

The GICC was supported by grants from the National Institutes of Health, Bethesda, Maryland (R01CA139020, R01CA52689, P50097257, P30CA125123, P30CA008748, C. Thompson PI). Additional support was provided by the McNair Medical Institute and the Population Sciences Biorepository at Baylor College of Medicine.

In Sweden work was additionally supported by Acta Oncologica through the Royal Swedish Academy of Science (BM salary) and The Swedish Research council and Swedish Cancer foundation. We are grateful to the National clinical brain tumor group, all clinicians and research nurses throughout Sweden who identified all cases.

The UCSF Adult Glioma Study was supported by the National Institutes of Health (grant numbers R01CA52689, P50CA097257, R01CA126831, and R01CA139020), the Loglio Collective, the National Brain Tumor Foundation, the Stanley D. Lewis and Virginia S. Lewis Endowed Chair in Brain Tumor Research, the Robert Magnin Newman Endowed Chair in Neuro-oncology, and by donations from families and friends of John Berardi, Helen Glaser, Elvera Olsen, Raymond E. Cooper, and William Martinusen. This project also was supported by the National Center for Research Resources and the National Center for Advancing Translational Sciences, National Institutes of Health, through UCSF-CTSI Grant Number UL1 RR024131. Its contents are solely the responsibility of the authors and do not necessarily represent the official views of the NIH. The collection of cancer incidence data used in this study was supported by the California Department of Public Health as part of the statewide cancer reporting program mandated by California Health and Safety Code Section 103885; the National Cancer Institute’s Surveillance, Epidemiology and End Results Program under contract HHSN261201000140C awarded to the Cancer Prevention Institute of California, contract HHSN261201000035C awarded to the University of Southern California, and contract HHSN261201000034C awarded to the Public Health Institute; and the Centers for Disease Control and Prevention’s National Program of Cancer Registries, under agreement # U58DP003862-01 awarded to the California Department of Public Health. The ideas and opinions expressed herein are those of the author(s) and endorsement by the State of California Department of Public Health, the National Cancer Institute, and the Centers for Disease Control and Prevention or their Contractors and Subcontractors is not intended nor should be inferred. Other significant contributors for the UCSF Adult Glioma Study include: M Berger, P Bracci, S Chang, J Clarke, A Molinaro, A Perry, M Pezmecki, M Prados, I Smirnov, T Tihan, K Walsh, J Wiemels, S Zheng.

GliomaScan group comprised: Laura E. Beane Freeman, Stella Koutros, Demetrius Albanes, Kala Visvanathan, Victoria L. Stevens, Roger Henriksson, Dominique S. Michaud, Maria Feychting, Anders Ahlbom, Graham G. Giles Roger Milne, Roberta McKean-Cowdin, Loic Le Marchand, Meir Stampfer, Avima M. Ruder, Tania Carreon, Goran Hallmans, Anne Zeleniuch-Jacquotte, J. Michael Gaziano, Howard D. Sesso, Mark P. Purdue, Emily White, Ulrike Peters, Howard D. Sesso, Julie Buring.

UK10K data generation and access was organized by the UK10K consortium and funded by the Wellcome Trust.

We are grateful to all the patients and individuals for their participation and we would also like to thank the clinicians and other hospital staff, cancer registries and study staff in respective centers who contributed to the blood sample and data collection.

## Supporting information captions

Supplemental Table 1. Case-only odds ratios (OR), 95% confidence intervals (95% CI), and p values from meta-analysis and individual studies for rs11979158, rs55705857 and rs9841110 overall and by histology groupings.

Supplemental Table 2. Characteristics of individuals in The Cancer Genome Atlas, by study and sex.

Supplemental Table 3. Linkage disequilibrium measures, sex-stratified odds ratios, and 95% confidence intervals (95% CI), and p values from meta-analysis for marker SNPs selected within the Cancer Genome Atlas genotyping data.

Supplemental Table 4. Odds ratios and 95% confidence intervals for unweighted scores in all glioma, GBM, and non-GBM overall and by sex.

Supplemental Table 5. Info score, sex-specific odds ratios (OR), 95% confidence intervals (95% CI), and p values from meta-analysis and individual studies for rs11979158, rs55705857 and rs9841110 overall and by histology groupings.

Supplemental Table 6. Risk allele frequencies (RAF), for meta-analysis and individual studies for rs11979158, rs55705857 and rs9841110 overall and by histology groupings.

Supplemental Table 7. Info score, sex-specific odds ratios (OR), 95% confidence intervals (95% CI), and p values from meta-analysis and individual studies for rs11979158, rs55705857 and rs9841110 by specific non-GBM histologies.

Supplemental Figure 1. P values of SNPs between 48.8mb and 50mb on chromosome 3 in males for A) all glioma, B) GBM, and C) non-GBM, and in females for D) all glioma, E) GBM, and F) non-GBM

Supplemental Figure 2. Proportion of samples by glioma subtype (based on IDH1/2 mutation, 1p19q, and TERT mutation) in the TCGA GBM and LGG datasets by sex, overall and stratified by study

Supplemental Figure 3. Proportion of samples by pan-glioma methylation subgroups [23] in the TCGA GBM and LGG datasets by sex, overall and stratified by study

Supplemental Figure 4. Density of histology-specific unweighted risk score by sex and case/control status for A) URS in all glioma, B) URS in GBM, C) URS in non-GBM, D) URS-GBM in GBM, only and E) URS-NGBM in non-GBM only

Supplemental Figure 5. Sex-specific odds ratios and 95% CI from meta-analysis and by study for rs11979158 (7p11.2) for all glioma, GBM, and non-GBM

Supplemental Figure 6. Sex-specific odds ratios and 95% CI from meta-analysis and by study for rs55705857 (8q24.21) for all glioma, GBM, and non-GBM

Supplemental Figure 7. Sex-specific odds ratios and 95% CI from meta-analysis and by study for rs9841110 (3p21.31) for all glioma, GBM, and non-GBM

## References

1. Ostrom QT, Gittleman H, Xu J, Kromer C, Wolinsky Y, et al. (2016) CBTRUS Statistical Report: Primary Brain and Other Central Nervous System Tumors Diagnosed in the United States in 2009–2013. Neuro-oncology 18: v1–v75.

2. Ostrom QT, Bauchet L, Davis F, Deltour I, Eastman C, et al. (2014) The epidemiology of glioma in adults: a “state of the science” review. Neuro-oncology 16: 896-913.

3. Kinnersley B, Mitchell JS, Gousias K, Schramm J, Idbaih A, et al. (2015) Quantifying the heritability of glioma using genome-wide complex trait analysis. Sci Rep 5: 17267.

4. Melin BS, Barnholtz-Sloan JS, Wrensch MR, Johansen C, Il’yasova D, et al. (2017) Genome-wide association study of glioma subtypes identifies specific differences in genetic susceptibility to glioblastoma and non-glioblastoma tumors. Nat Genet 49: 789-794.

5. Benson VS, Kirichek O, Beral V, Green J (2015) Menopausal hormone therapy and central nervous system tumor risk: large UK prospective study and meta-analysis. Int J Cancer 136: 2369-2377.

6. Zong H, Xu H, Geng Z, Ma C, Ming X, et al. (2014) Reproductive factors in relation to risk of brain tumors in women: an updated meta-analysis of 27 independent studies. Tumour Biol 35: 11579-11586.

7. Howlader N NA, Krapcho M, Miller D, Bishop K, Kosary CL, Yu M, Ruhl J, Tatalovich Z, Mariotto A, Lewis DR, Chen HS, Feuer EJ, Cronin KA (eds) (2017) SEER Cancer Statistics Review, 1975-2014, based on November 2016 SEER data submission. Bethesda, MD: National Cancer Institute.

8. Siegel RL, Miller KD, Jemal A (2017) Cancer Statistics, 2017. CA Cancer J Clin 67: 7-30.

9. Liu LY, Schaub MA, Sirota M, Butte AJ (2012) Sex differences in disease risk from reported genome-wide association study findings. Hum Genet 131: 353-364.

10. Dorak MT, Karpuzoglu E (2012) Gender differences in cancer susceptibility: an inadequately addressed issue. Front Genet 3: 268.

11. Amirian ES, Armstrong GN, Zhou R, Lau CC, Claus EB, et al. (2016) The Glioma International Case-Control Study: A Report From the Genetic Epidemiology of Glioma International Consortium. Am J Epidemiol 183: 85-91.

12. Wrensch M, Jenkins RB, Chang JS, Yeh RF, Xiao Y, et al. (2009) Variants in the CDKN2B and RTEL1 regions are associated with high-grade glioma susceptibility. Nature Genetics 41: 905-908.

13. Shete S, Hosking FJ, Robertson LB, Dobbins SE, Sanson M, et al. (2009) Genome-wide association study identifies five susceptibility loci for glioma. Nature Genetics 41: 899-904.

14. Rajaraman P, Melin BS, Wang Z, McKean-Cowdin R, Michaud DS, et al. (2012) Genome-wide association study of glioma and meta-analysis. Human Genetics 131: 1877-1888.

15. Yeager M, Chatterjee N, Ciampa J, Jacobs KB, Gonzalez-Bosquet J, et al. (2009) Identification of a new prostate cancer susceptibility locus on chromosome 8q24. Nat Genet 41: 1055-1057.

16. Hunter DJ, Kraft P, Jacobs KB, Cox DG, Yeager M, et al. (2007) A genome-wide association study identifies alleles in FGFR2 associated with risk of sporadic postmenopausal breast cancer. Nat Genet 39: 870-874.

17. Sahm F, Reuss D, Koelsche C, Capper D, Schittenhelm J, et al. (2014) Farewell to oligoastrocytoma: in situ molecular genetics favor classification as either oligodendroglioma or astrocytoma. Acta Neuropathol 128: 551-559.

18. Machiela MJ, Chanock SJ (2015) LDlink: a web-based application for exploring population-specific haplotype structure and linking correlated alleles of possible functional variants. Bioinformatics 31: 3555-3557.

19. Sanson M, Hosking FJ, Shete S, Zelenika D, Dobbins SE, et al. (2011) Chromosome 7p11.2 (EGFR) variation influences glioma risk. Hum Mol Genet 20: 2897-2904.

20. Filardo EJ (2002) Epidermal growth factor receptor (EGFR) transactivation by estrogen via the G-protein-coupled receptor, GPR30: a novel signaling pathway with potential significance for breast cancer. J Steroid Biochem Mol Biol 80: 231-238.

21. Sun T, Warrington NM, Luo J, Brooks MD, Dahiya S, et al. (2014) Sexually dimorphic RB inactivation underlies mesenchymal glioblastoma prevalence in males. J Clin Invest 124: 4123-4133.

22. Enciso-Mora V, Hosking FJ, Kinnersley B, Wang Y, Shete S, et al. (2013) Deciphering the 8q24.21 association for glioma. Hum Mol Genet 22: 2293-2302.

23. Jenkins RB, Xiao Y, Sicotte H, Decker PA, Kollmeyer TM, et al. (2012) A low-frequency variant at 8q24.21 is strongly associated with risk of oligodendroglial tumors and astrocytomas with IDH1 or IDH2 mutation. Nat Genet 44: 1122-1125.

24. Ceccarelli M, Barthel FP, Malta TM, Sabedot TS, Salama SR, et al. (2016) Molecular Profiling Reveals Biologically Discrete Subsets and Pathways of Progression in Diffuse Glioma. Cell 164: 550-563.

25. Brennan CW, Verhaak RG, McKenna A, Campos B, Noushmehr H, et al. (2013) The somatic genomic landscape of glioblastoma. Cell 155: 462-477.

26. The Cancer Genome Atlas Research Network, Brat DJ, Verhaak RG, Aldape KD, Yung WK, et al. (2015) Comprehensive, Integrative Genomic Analysis of Diffuse Lower-Grade Gliomas. N Engl J Med 372: 2481-2498.

27. Consortium GT (2013) The Genotype-Tissue Expression (GTEx) project. Nat Genet 45: 580-585.

28. Broad Institute TCGA Genome Data Analysis Center (2016) Analysis-ready standardized TCGA data from Broad GDAC Firehose 2016 01 28 run. Broad Institute of MIT and Harvard.

29. Hayashi Y, Yamaguchi J, Kokuryo T, Ebata T, Yokoyama Y, et al. (2016) The Complete Loss of Tyrosine Kinase Receptors MET and RON Is a Poor Prognostic Factor in Patients with Extrahepatic Cholangiocarcinoma. Anticancer Res 36: 6585-6592.

30. Lee JR, Roh JL, Lee SM, Park Y, Cho KJ, et al. (2017) Overexpression of glutathione peroxidase 1 predicts poor prognosis in oral squamous cell carcinoma. J Cancer Res Clin Oncol 143: 2257-2265.

31. Elks CE, Perry JR, Sulem P, Chasman DI, Franceschini N, et al. (2010) Thirty new loci for age at menarche identified by a meta-analysis of genome-wide association studies. Nat Genet 42: 1077-1085.

32. Perry JR, Day F, Elks CE, Sulem P, Thompson DJ, et al. (2014) Parent-of-origin-specific allelic associations among 106 genomic loci for age at menarche. Nature 514: 92-97.

33. Pokrajac-Bulian A, Toncic M, Anic P (2015) Assessing the factor structure of the Body Uneasiness Test (BUT) in an overweight and obese Croatian non-clinical sample. Eat Weight Disord 20: 215-222.

34. Raelson JV, Little RD, Ruether A, Fournier H, Paquin B, et al. (2007) Genome-wide association study for Crohn’s disease in the Quebec Founder Population identifies multiple validated disease loci. Proc Natl Acad Sci U S A 104: 14747-14752.

35. Reinius B, Saetre P, Leonard JA, Blekhman R, Merino-Martinez R, et al. (2008) An evolutionarily conserved sexual signature in the primate brain. PLoS Genet 4: e1000100.

36. Rinn JL, Snyder M (2005) Sexual dimorphism in mammalian gene expression. Trends Genet 21: 298-305.

37. Ellegren H, Parsch J (2007) The evolution of sex-biased genes and sex-biased gene expression. Nat Rev Genet 8: 689-698.

38. Sun T, Plutynski A, Ward S, Rubin JB (2015) An integrative view on sex differences in brain tumors. Cell Mol Life Sci 72: 3323-3342.

39. Lichtenstein P, Holm NV, Verkasalo PK, Iliadou A, Kaprio J, et al. (2000) Environmental and heritable factors in the causation of cancer--analyses of cohorts of twins from Sweden, Denmark, and Finland. N Engl J Med 343: 78-85.

40. Ahmed S, Thomas G, Ghoussaini M, Healey CS, Humphreys MK, et al. (2009) Newly discovered breast cancer susceptibility loci on 3p24 and 17q23.2. Nat Genet 41: 585-590.

41. Amos CI, Dennis J, Wang Z, Byun J, Schumacher FR, et al. (2016) The OncoArray Consortium: a Network for Understanding the Genetic Architecture of Common Cancers. Cancer Epidemiol Biomarkers Prev.

42. Li Y, Byun J, Cai G, Xiao X, Han Y, et al. (2016) FastPop: a rapid principal component derived method to infer intercontinental ancestry using genetic data. BMC Bioinformatics 17: 122.

43. Delaneau O, Marchini J, Zagury JF (2012) A linear complexity phasing method for thousands of genomes. Nat Methods 9: 179-181.

44. Howie BN, Donnelly P, Marchini J (2009) A flexible and accurate genotype imputation method for the next generation of genome-wide association studies. PLoS Genet 5: e1000529.

45. Genomes Project C, Auton A, Brooks LD, Durbin RM, Garrison EP, et al. (2015) A global reference for human genetic variation. Nature 526: 68-74.

46. Golding J, Pembrey M, Jones R, Team AS (2001) ALSPAC--the Avon Longitudinal Study of Parents and Children. I. Study methodology. Paediatr Perinat Epidemiol 15: 74-87.

47. Moayyeri A, Hammond CJ, Hart DJ, Spector TD (2013) The UK Adult Twin Registry (TwinsUK Resource). Twin Res Hum Genet 16: 144-149.

48. Purcell S, Chang C PLINK 1.9.

49. Das S, Forer L, Schonherr S, Sidore C, Locke AE, et al. (2016) Next-generation genotype imputation service and methods. Nat Genet 48: 1284-1287.

50. Loh PR, Danecek P, Palamara PF, Fuchsberger C, Y AR, et al. (2016) Reference-based phasing using the Haplotype Reference Consortium panel. Nat Genet 48: 1443-1448.

51. McCarthy S, Das S, Kretzschmar W, Delaneau O, Wood AR, et al. (2016) A reference panel of 64,976 haplotypes for genotype imputation. Nat Genet 48: 1279-1283.

52. Eckel-Passow JE, Lachance DH, Molinaro AM, Walsh KM, Decker PA, et al. (2015) Glioma Groups Based on 1p/19q, IDH, and TERT Promoter Mutations in Tumors. N Engl J Med 372: 2499-2508.

53. Marchini J, Howie B, Myers S, McVean G, Donnelly P (2007) A new multipoint method for genome-wide association studies by imputation of genotypes. Nat Genet 39: 906-913.

54. Paternoster R, Brame R, Mazerolle P, Piquero A (1998) USING THE CORRECT STATISTICAL TEST FOR THE EQUALITY OF REGRESSION COEFFICIENTS. Criminology 36: 859-866.

55. Mittelstrass K, Ried JS, Yu Z, Krumsiek J, Gieger C, et al. (2011) Discovery of Sexual Dimorphisms in Metabolic and Genetic Biomarkers. PLOS Genetics 7: e1002215.

56. Liu JZ, Tozzi F, Waterworth DM, Pillai SG, Muglia P, et al. (2010) Meta-analysis and imputation refines the association of 15q25 with smoking quantity. Nat Genet 42: 436-440.

57. Chow JC, Yen Z, Ziesche SM, Brown CJ (2005) Silencing of the mammalian X chromosome. Annu Rev Genomics Hum Genet 6: 69-92.

58. R Core Team (2017) R: A language and environment for statistical computing. Vienna, Austria: R Foundation for Statistical Computing.

59. Wickham H (2009) ggplot2: elegant graphics for data analysis.: Springer New York.

60. Karssen LC, van Duijn CM, Aulchenko YS (2016) The GenABEL Project for statistical genomics. F1000Res 5: 914.

61. Turner SD (2014) qqman: an R package for visualizing GWAS results using Q-Q and manhattan plots. biorXiv DOI: 10.1101/005165.

62. Mittlbock M, Schemper M (1996) Explained variation for logistic regression. Stat Med 15: 1987-1997.

